# A termination-aware vector design improves heterologous gene expression in *Pseudomonas putida*

**DOI:** 10.1101/2022.11.22.517491

**Authors:** Guilherme Marcelino Viana de Siqueira, María-Eugenia Guazzaroni

## Abstract

Transcriptional terminators are key players in the flow of genetic information, but are often overlooked in circuit design. In this work, we used the Standard European Vector Architecture (SEVA) as a scaffold to investigate the effects of different terminators in the output of a reporter construct expressed in two bacterial species and found that replacing the conventional T1 and T0 transcriptional terminators of the SEVA vector format with a set of broad-host metagenomic terminators resulted in a significant improvement in the signal of a fluorescent device in *Pseudomonas putida* KT2440. Interestingly, this new vector design displayed the same performance as pSEVA231 in *Escherichia coli* DH10B. Our results support the notion that transcriptional terminators may affect the interconnected mesh of biological processes taking place in gene expression, leading to a host-dependent output that should be heeded for optimal circuit expression in different microbial hosts.

## Introduction

First published just shy of a decade ago, the SEVA family of vectors was conceived as a standardized plasmid platform that aimed to simplify the portability of genetic circuits to a wide range of Gram-negative microbial hosts^1^. Due to this attractive premise, the SEVA initiative has garnered the attention of a wide community of synthetic biologists worldwide that has since helped expand upon this initial concept by spawning several derivatives, many of which have been acknowledged in subsequent revisions of the standard and incorporated into the SEVA database^2,3^. Still, even after many iterations, canonical SEVA plasmids and the so-called SEVA siblings majorly rely on a single pair of transcriptional terminators for insulating most DNA constructions placed in the cargo module of any given vector, namely the 105 bp T1 terminator of the *E. coli rrnB* operon and the 103 bp T0 terminator from the phage lambda^1^.

Despite acting as structural elements of the SEVA format, transcriptional terminators have been shown to possess context- and host-dependent efficiency^4,5^ that may lead to unexpected consequences in processes such as the rate of plasmid replication^6^ and the expression of extra-construct genes^7^. Given the importance of these genetic parts for a proper expression of synthetic constructs, in this work, we took upon investigating the effects of replacing the standard terminators of a SEVA plasmid in the expression of a simple reporter device by two microbial hosts. While this approach may be further streamlined in future efforts, here we revisit the Standard European Vector Architecture and provide a blueprint for introducing new termination sequences to conventional SEVA backbones, which can be leveraged for fine-tuning the performance of synthetic circuits in non-conventional bacteria.

## Results and Discussion

In this work, we devised a new cargo architecture for SEVA plasmids, including novel transcriptional terminators flanking both ends of a standard Multiple Cloning Site (MCS). Since this design does not readily comply with the canonical SEVA nomenclature, we opted for naming the resulting module VANT (*Variant Architecture with New Terminators*), and, for simplicity, refer to the modified pSEVA231 vector described here as pVANT. The fundamental feature of the 797 bp-long VANT module is the presence of four environmental transcriptional terminators (eTs) insulating the MCS. These genetic parts were chosen based on a previous functional screening of a soil metagenomic library and have shown high efficiency in several Proteobacteria^8^. Within the module, the four eTs are distributed in two groups placed at the 5’ and 3’ termini of the MCS, each containing a terminator in the sense and antisense transcription directions to account for readthrough from within the MCS and from the plasmid backbone (**Figure 1**). Despite these design choices, the VANT module follows the conventions established in the SEVA standard, i.e. it is flanked by the restriction sites *Asc*I and *Swa*I (as the T1 and T0 terminators originally are) and it does not include reserved restriction sites outside of the MCS^1^. These features make the module readily transposable to any SEVA backbone by routine cloning and assembly techniques. A description of useful overlapping sites and primer designs for the introduction of the VANT module in standard SEVA backbones is provided in the supplementary material (**Figure S1, Table S1**).

**Figure 1.**
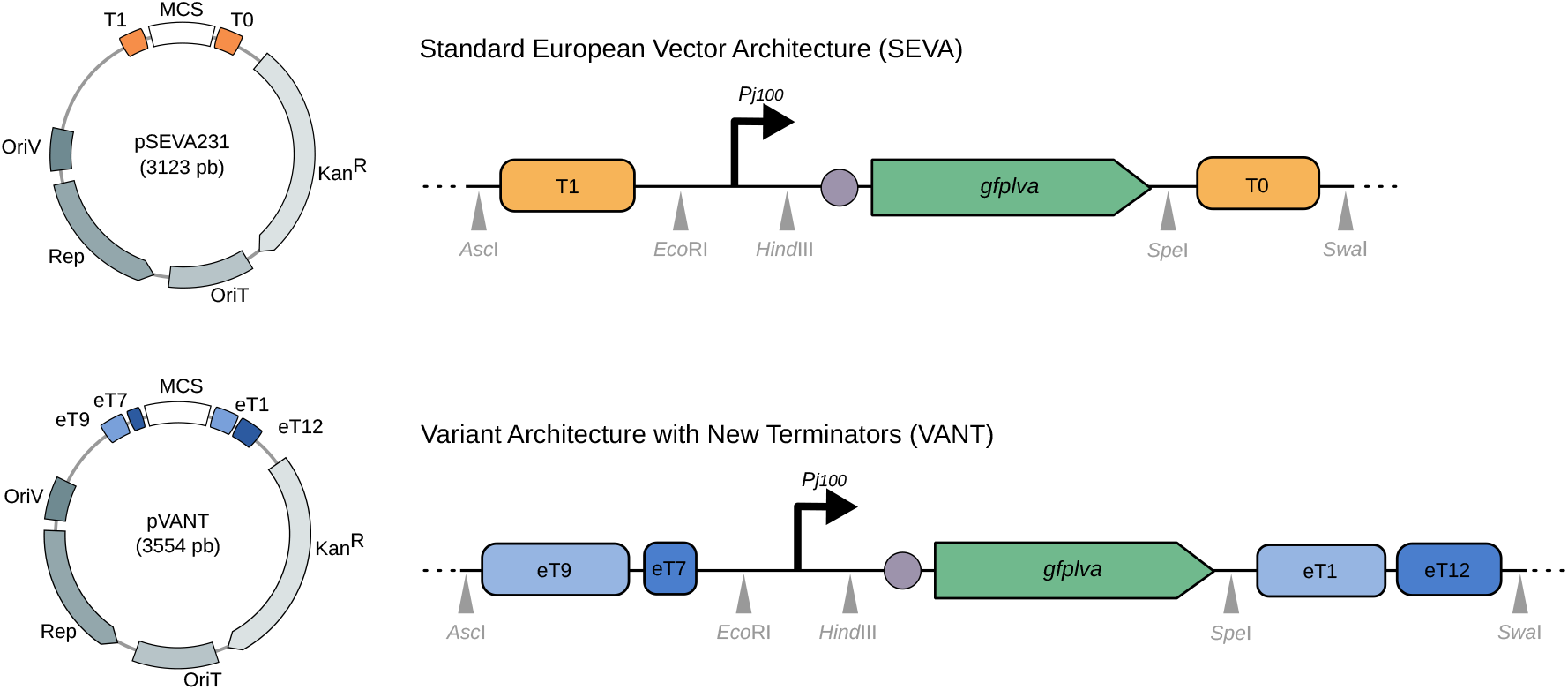
**A)** Illustrative representation of vectors pSEVA231 (top) and pVANT (bottom). These vectors share several structures, such as the MCS, the antibiotic selection marker Kan^R^, the origin of transfer OriT, and the pBBR1-based replication origin OriV with its associated *rep* genes (shown in different shades of gray), but differ regarding the transcriptional terminators flanking the MCS (highlighted in blue/orange). In this representation, the VANT terminators colored in light blue are placed in the sense direction (considering 5’ → 3’ as left to right) and darker blue terminators are placed in the antisense direction. In the pSEVA, the T1 and T0 terminators are in the sense direction^1^. **B)** Detailed view of the MCS of pSEVA231/pVANT vectors. In both vectors, the reporter protein GFP*lva* and a RBS sequence (gray circle) were cloned between the *Eco*RI and *HindIII* restriction sites and introduced into *E. coli* DH10B and *P. putida* KT2440. For each vector, *gfplva* was cloned by itself or under control of a strong constitutive promoter (*Pj100*).

As a proof-of-concept for this this new vector design, we evaluated the performance of pVANT and pSEVA231 in both the model environmental bacterium *Pseudomonas putida* KT2440 and the conventional laboratory strain *Escherichia coli* DH10B by measuring fluorescence of the GFP*lva* reporter protein under regulation of a constitutive synthetic promoter both at the population and single-cell levels (**Figure 2**). Our results show that, under the tested conditions, the use of either vector architecture did not affect the measured expression levels of the reporter in *E. coli*. In fact, in microplate cultivations the growth profiles of *E. coli* cells harboring any of the vectors was fairly constant across all experiments and the observed expression levels of the fluorescent reporter were similar in intermediate points of the curve (**Figure 2A**), even though the overall expression profile throughout the growth curves appeared to be more evened out with pVANT than with pSEVA231 (**Figure S2**). On the other hand, it is evident that, in *P. putida* KT2440, the measured fluorescence levels differed significantly depending on the vector tested, with pVANT outperforming its SEVA counterpart (**Figure 2A**). Single-cell level assays suggest that this overall weaker fluorescence signal observed for *pSEVA231-Pj100-gfplva* in plate reader experiments may be caused by the formation of a considerable subpopulation of non-fluorescent cells (**Figure 2B**), which may indicate poorer plasmid replication for this particular strain under the tested conditions^9^. Conversely, results for *E. coli* DH10B are consistent between pSEVA231 and pVANT across microplate reader and cytometry experiments. This agrees with previous results showing that termination efficiency is less dependent on terminator sequence in *E. coli* than in other bacteria^8^. Put together, our data indicates that, in comparison to a conventional SEVA vector, the sort of intervention we propose can result in a positive impact in the output of a reporter device in *P. putida* KT2440 while having negligible effects in a conventional laboratory strain of *E. coli*.

**Figure 2.**
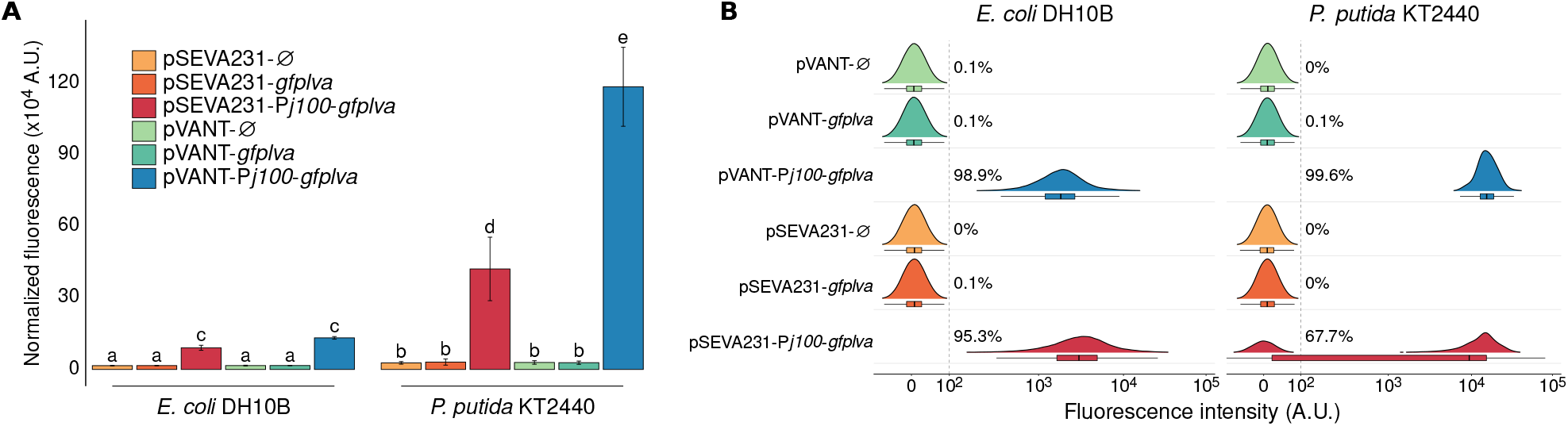
**A)** Normalized fluorescence measured for cell populations at the fourth hour of incubation in microplates. Two-way ANOVA revealed a statistically significant interaction between the plasmid architecture and host choice on normalized fluorescence levels (F(5, 45) = 27.11, p = 1.54e-12). Different significance groups, based on a Tukey *post-hoc* test, are reported above the bars. **B)** Representative single-cell fluorescence profiles for bacteria carrying pSEVA231 or pVANT plasmids. The dotted line represents an arbitrary threshold to define fluorescent and non-fluorescent populations. The percentages refer to the proportion of fluorescent cells in each sample. A.U. = arbitrary units.

Standardization and reliability of genetic parts have always been among the greatest efforts in Synthetic Biology, and even though great strides have been made in setting the groundwork for this discipline, the introduction of different microbial hosts adds a new layer of complexity on top of an already challenging task^10,11^. This has recently been addressed as “contextual dependency” in a series of works focused on the implementation of genetic inverters (NOT gates), originally characterized in *E. coli*, for automating the design of complex logical gates in *P. putida^12–14^*. Considering the relatively underappreciated effects of transcriptional terminators for the contextual dependency of synthetic constructs, we believe that the kind of plasmid refactoring proposed in this work may be useful for optimizing and fine-tuning the performance of genetic devices intended to be used across different microbial hosts, aiding in the introduction of more complex behaviors in non-conventional bacteria.

## Methods

### Design of the VANT module and construction of the reporter vectors

The GenBank sequence for pSEVA231 (accession JX560328) was used as the template for the design of the new VANT module. We used the Unipro UGENE toolkit (version 39)^15^ for replacing the terminator sequences found in the original plasmid with those described in Amarelle et al^8^ as mentioned in the text. The entire pVANT sequence was synthesized *de novo* (GenScript, NJ, USA) and cloned into *E. coli* DH10B before being manipulated in the subsequent steps of this work. To construct the reporter vectors, the *gfplva^16^* gene was cloned between the *HindIII* and *SpeI* restriction sites both in pSEVA231 and pVANT vectors. After this, each vector containing *gfplva* was amplified with the primer pairs P5/P8 and P7/P6 (**Figure S1**, **Table S1**) for introducing the *Pj100* promoter sequence into the plasmid via Gibson assembly. Confirmation of assembly was performed by the amplification of construct-specific plasmid regions, by restriction analysis, and visually by the fluorescence phenotype of plated cells.

### Microplate and cytometry assays

*E. coli* DH10B or *P. putida* KT2440 carrying different pVANT/pSEVA231 vectors were inoculated into M9 media (composed of M9 salts: 7,5 g/L Na_2_HPO_4_·H_2_O, 3 g/L KH_2_PO_4_, 1 g/L NH_4_Cl, 0.5 g/L NaCl, with the addition of 2mM MgSO_4_ and 0.1mM CaCl_2_) supplemented with 1% glycerol (*E. coli*) or 1% citrate (*P. putida*) and 0.1% cas-amino acids, and incubated overnight at 37°C (*E. coli*) or 30°C (*P. putida*), with agitation at 220 rpm. For all cultivations 50 μg/mL of kanamicin was used for selection of the plasmids. Following incubation, each culture was diluted to 0.05 OD_600_ in fresh M9 media with the appropriate carbon source for each microbial host. For the microplate reader assays, 200 μL of these dilutions were then pipetted into opaque 96-well microplates and incubated at 30°C in a Victor X3 multilabel microplate reader (Perkin Elmer) that periodically measured absorbance (OD_600_) and fluorescence (Ex: 488 nm / Em: 535 nm) for eight hours. For the flow cytometry assays, the saturated overnight cultures were diluted to 0.05 OD in 5 mL of fresh media and cultivated under the described conditions for four additional hours. After this, the tubes were placed in ice to halt cellular growth and cells were promptly loaded in a Canto II flow cytometer (BD). GFPlva was excited with a 488 nm blue laser and detected in the FITC-A channel with a 530/30 band pass filter. In each experiment, 100.000 events of each sample were recorded.

### Data Analysis

R version 4.1.2 was used for all data processing and statistical analyses performed in this work. Flow cytometry data, exported as .fcs files, was processed using the flowCore (version 2.8.0)^17^ and flowStats (version 4.8.2)^18^ packages. Data exported from the microplate reader was parsed and imported into R using the mipreadr package version 0.1.0 (github.com/viana-guilherme/mipreadr). In addition to these, the tidyverse suite of packages (version 1.3.2) was extensively used for manipulating and visualizing data. A more comprehensive listing of the packages used, as well as raw data from each experiment and reproducible code examples are provided in the supplementary material.

## Author Information

### Notes

The authors declare no competing financial interest.

## Acknowledgments

The authors would like to thank the lab technicians Thalita Riul Prado and Denise Brufato Ferraz for their invaluable assistance in the course of this work. We are also thankful to our lab colleagues for the insightful discussions during the preparation of this manuscript. This work was supported by the São Paulo Research Foundation (FAPESP) (awards # 2019/25432-7 and # 2021/01748-5).

## Notes

### Competing Interest Statement

The authors have declared no competing interest.

