## Supplemental Figure S1, Figure S2 and Table S1 for "A termination-aware vector design improves heterologous gene expression in *Pseudomonas putida*"

**by**

Guilherme Marcelino Viana de Siqueira[1] and María-Eugenia Guazzaroni[1*]

[1] Department of Biology, Faculty of Philosophy, Sciences and Letters at Ribeirão Preto, University of São Paulo, Brazil

### Supporting Information

#### *Abstract*

Transcriptional terminators are key players in the flow of genetic information, but are often overlooked in circuit design. In this work, we used the Standard European Vector Architecture (SEVA) as a scaffold to investigate the effects of different terminators in the output of a reporter construct expressed in two bacterial species and found that replacing the conventional T1 and T0 transcriptional terminators of the SEVA vector format with a set of broad-host metagenomic terminators resulted in a significant improvement in the signal of a fluorescent device in *Pseudomonas putida* KT2440. Interestingly, this new vector design displayed the same performance as pSEVA231 in *Escherichia coli* DH10B. Our results support the notion that transcriptional terminators may affect the interconnected mesh of biological processes taking place in gene expression, leading to a host-dependent output that should be heeded for optimal circuit expression in different microbial hosts.

*Please note that raw data and code used in this work are available upon request.*

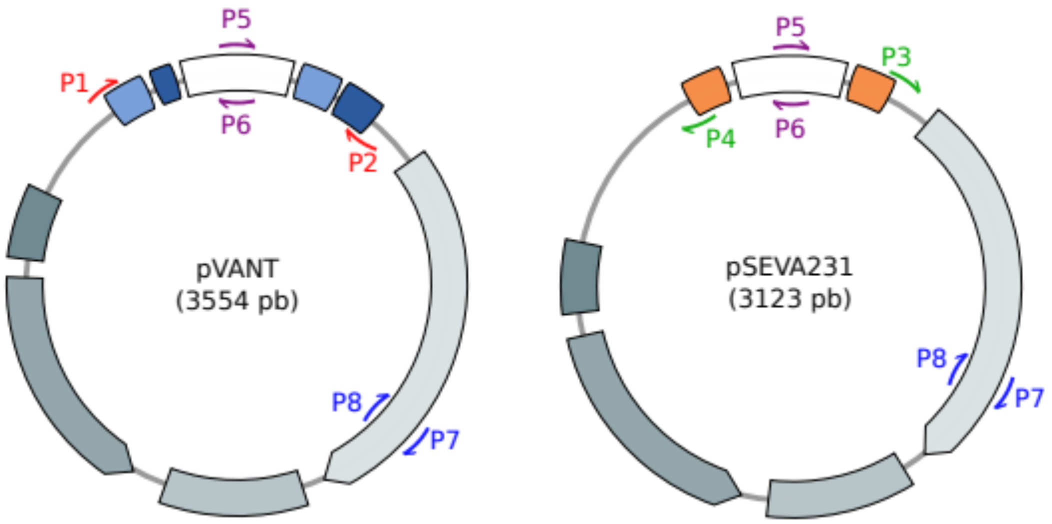


**Figure S1.** Illustrative representation of the relative positions of binding sites for the primers described in **Table S1** in their context within the sequence of vectors pVANT and pSEVA231 (not to scale).

**Table S1.** Primers used in this study.

| ID | Sequence (5' → 3') | Length (nt) | Usage notes |
|---|---|---|---|
| P1 | CAGCTGTCTAGGAGATAGCTAC | 22 | These primers are used for the amplification of the VANT module. Short sequences of the original T1 and T0 terminators remain in the amplicon, helping constitute overlapping regions with the SEVA backbones for Gibson Assembly |
| P2 | CCTGGATTCTCACCTCGTAC | 20 | |
| P3 | GCACTACATTGTACGAGGTGAGAAT CCAGGGGTCC | 35 | These primers anneal to the terminators on each end of the SEVA backbone and introduce overlapping regions with the VANT module. After pVANT is assembled, they can be used to amplify the backbone directly from it as a template. |
| P4 | GCCATGTTGTAGCTATCTCCTAGACA GCTGGGCGC | 35 | |
| P5 | **TTGACGGCTAGCTCAGTCCTAGGTA CAGTGCTAGC**AAGTGTCGACCTGCA GGCATGC | 57 | These primers introduce the constitutive promoter BBa_J23100 from the iGEM collection[1] (referred to as P*j100* in the text, and shown here in bold) as overhangs in the amplicons. In relation to the original sequence, we added a small 3' spacer (AAGT) after the promoter sequence. Restriction sites of the SEVA MCS are used as binding sites for PCR amplification. The promoter sequence itself acts as the overlapping region for Gibson assembly. |
| P6 | ACTT**GCTAGCACTGTACCTAGGACT GAGCTAGCCGTCAA**GGGTACCGAG CTCGAATTC | 58 | |

1 http://parts.igem.org/Promoters/Catalog/Anderson

| | | | |
|---|---|---|---|
| P7 | GCGGATCGTTATCAGGATCTG | 21 | These primers anneal to the antibiotic resistance marker in the plasmid backbone. They are designed so as to form a 40 nt overlap in between their amplicons, which may be useful for assembly strategies that involve splitting the backbone into separate fragments. |
| P8 | GGCAGTTCCACAGAATGGC | 19 | |

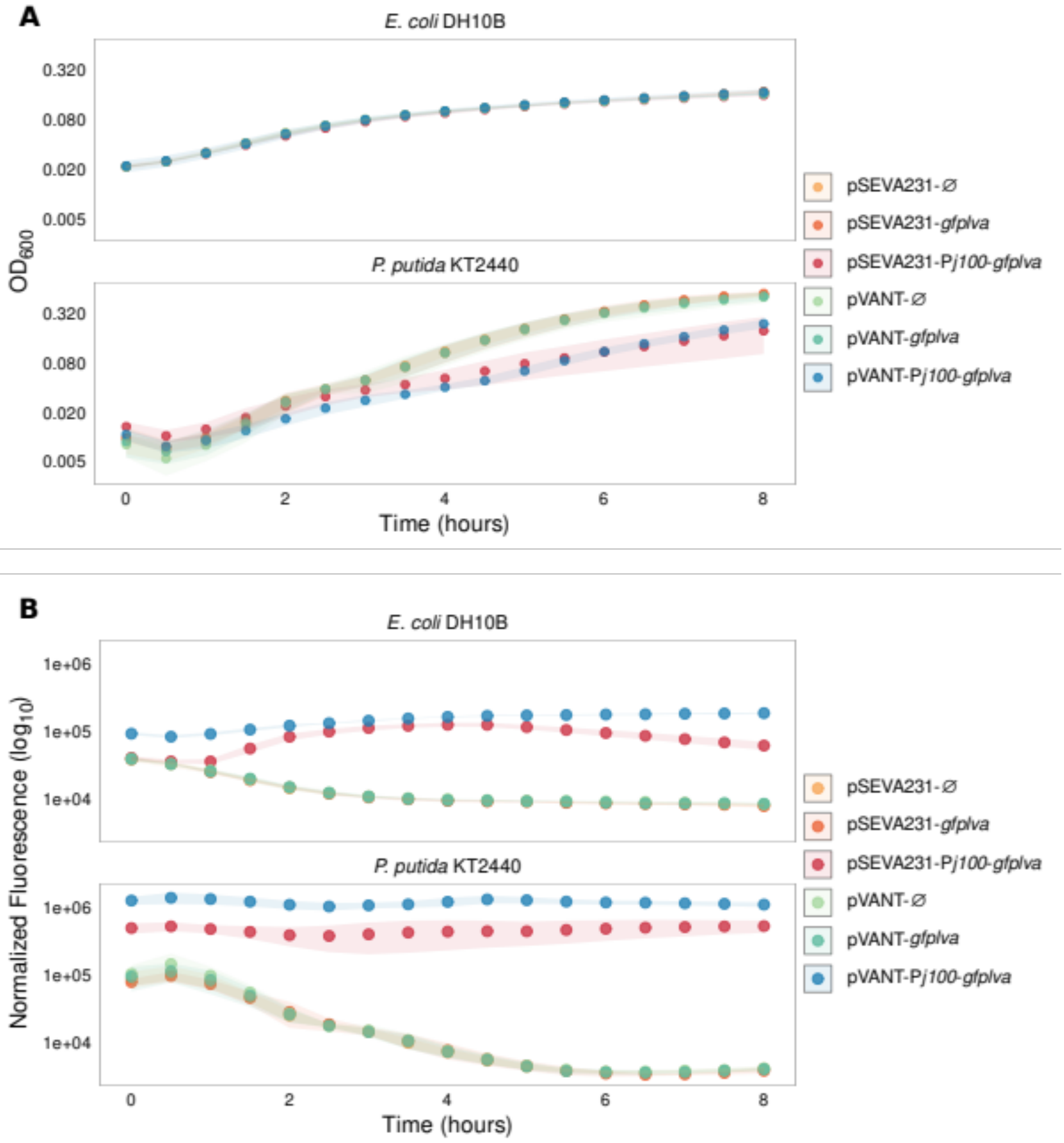


**Figure S2. A.** Growth curves of *E. coli* DH10B (top) and *P. putida* KT2440 (bottom) in 96-well microplates. In the figure, plotted in a semi-logarithmic scale, the dots represent the mean and the shaded areas, the standard deviations of at least four biological replicates. **B.** Expression profiles of GFP*lva* in *E. coli* DH10B (top) and *P.*

*putida* KT2440 (bottom). Fluorescence levels measured by the microplate reader were normalized by the optical density of growing cells.
